# Lab-on-a-graphene-FET detection of key molecular events underpinning influenza virus infection and effect of antiviral drugs

**DOI:** 10.1101/2020.03.18.996884

**Authors:** T. Ono, K. Kamada, R. Hayashi, A. R. Piacenti, C. Gabbutt, N. Sriwilaijaroen, H. Hiramatsu, Y. Kanai, K. Inoue, S. Nakakita, T. Kawahara, Y. Ie, Y. Watanabe, Y. Suzuki, S. Contera, K. Matsumoto

## Abstract

Small solid-state devices are candidates for accelerating biomedical assays/drug discovery, however their potential remains unfulfilled. Here, we demonstrate that graphene-field effect transistors (FET) can be used to successfully detect the key molecular events underlying viral infections and the effect of antiviral drugs. Our device success is achieved by bio-mimicking the host-cell surface during an influenza infection at the graphene channel. In-situ AFM confirms the biological interactions at the sialic acid-functionalized graphene: viral hemagglutinin (HA) binds to sialic acid, and neuraminidase (NA) reacts with the sialic acid-HA complex. The graphene-FET detects HA binding to sialic acid, and NA cleavage of sialic acid. The inhibitory effect of the drug “zanamivir” on NA-sialic acid interactions is monitored in real-time; the reaction rate constant of NA-sialic acid reaction was successfully determined. We demonstrate that graphene-FETs are powerful platforms for measurement of biomolecular interactions and contribute to future deployment of solid-state devices in drug discovery/biosensing.

## Introduction

Influenza virus is amongst the most deadly human pathogens. The main reason for its virulence is the emergence of pandemics by the evolution of particularly lethal influenza viral strains. The standard pattern of an influenza infection in adults is characterized by an exponential growth of virus titer, which peaks 2 to 3 days post-infection, therefore infections can spread very quickly. Furthermore influenza achieves a very efficient airborn transmissibility (*1*). At the molecular level, the success of an influenza infection is determined by two key events on the cell surface: (i) the initial binding of the infecting virus to the host cell, and (ii) the detachment of the newly made viruses from surface of the host cell after replication. Both events are mediated by sialic acid present at carbohydrate side chains of glycoproteins and glycolipids at the surface of host cells. Infection initiates with the binding of hemagglutinin (HA) proteins present at the virus surface to surface sialic acid groups (*2*). Upon replication, the enzyme neuraminidase (NA) binds to and cleaves the sialic acid site of the glyco-biomolecules at the surfaces of infected cell; upon cleavage, viruses produced in the infected cell are released (*3*). Both events are the targets of antibodies that confer protective immunity to influenza viruses (*4–6*) and are the most effective pharmacological antiviral targets. The three influenza antiviral drugs currently approved by Food and Drug Administration of the USA — peramivir (trade name Rapivab™), oseltamivir phosphate (Tamiflu™) and zanimivir (Relenza™)—are NA inhibitors. Anti-influenza drugs mimic the natural sialic acid substrate of NA but achieve a much tighter molecular binding improve efficacy (*2*). Influenza viruses can become resistant to influenza antiviral drugs, and hence fast methods of identifying key molecular interactions and facilitation of antiviral drug discovery are fundamental to combat future pandemics.

Drug discovery typically investigates interactions between a lead compound (e.g. a potential drug) and a target (e.g. a protein or a molecular interaction). Drug discovery requires the testing of millions of different chemical combinations, hence high-throughput systems for handling large number of samples must be able to work in parallel and utilize small chemical volumes to keep the cost affordable (*7*). In this context, small “lab-on-a chip” devices are emerging as important future players in the field (*8*).

In particular the development of biomimetic device-platforms that make possible the in-situ monitoring of the molecular details of key viral infection activities and the effect of drugs, can greatly accelerate and reduce costs of research on antiviral drugs. The sensitivity and surface-based measurements of graphene make it a very attractive material to achieve this goal. The extraordinarily high electron mobility of graphene makes it highly sensitive to changes in the surrounding environment (*9*), and hence graphene is considered an excellent material candidate for detecting interactions and reactions in very small samples of biomolecules immobilized on its surface by acting as channel in field effect transistors (FET) in high sensitivity biosensor applications (*10*). Furthermore, the large surface, good electronic conductivity, wide electrochemical window, high stability, sensitivity, low detection limit facilitate the application of electric fields for bio-detection in aqueous solution (*10–12*). As graphene simultaneously plays the role of a sensing surface and the conducting channel, its sensitivity can potentially be very high (*9*). It has been demonstrated that solution-gated graphene-FETs can work as very effective pH and protein concentration sensors (*11,13*) and have demonstrated their capacity for label-free detection of peptide nucleic acid-DNA hybridization (*14*) and unamplified target genes (*15*). Furthermore, the 2-dimensional nature of graphene and the ease of chemical modification and molecular immobilization (e.g. by using linker molecules able to form π-π stacking interactions at its surface (*16,17*)) make it a useful substrate for uniform and controlled functionalization of biomolecules in planar geometries (*18*) that can be designed to mimic e.g. the disposition of proteins at biological surfaces such as the cell membrane.

Here we propose to utilize a solution-gated graphene-FET as a “lab-on-a-graphene-FET” device to study the activity of HA and NA on a biomimetic graphene channel. We create a surface that mimics the membrane surface of the host cell to an influenza virus infection by immobilization of the key molecules at the graphene surface (sugar chains containing sialic acid groups). Furthermore we demonstrate that it is possible to monitor the activities of HA and NA and the effect of the drug zanamivir with the device.

The structural details of the molecular activity at the surface of the FET are investigated using atomic force microscopy (AFM) imaging in solution. The combination of high-resolution AFM imaging and FET current measurements allows us to characterize the binding of molecules at the surface of graphene, the structural effect of the enzymes and drugs on the substrate and to correlate structural changes at the graphene electrode with the read-out measurements of the FET device.

## Results and Discussion

### Preparation and characterization of a host-cell membrane biomimetic surface to function as channel in a graphene-FET

1-pyrenebutanoic acid succinimidyl ester (PBASE) molecules (see Materials and Methods) was immobilized on the graphene surface by π-π interactions. PBASE is used as a linker to subsequently functionalize the surface with sialoglycopeptide (SGP) sugar chains containing sialic acid. Figure 1A is a typical AFM image of the graphene with gold electrodes. Uniform monolayer graphene was identified in aqueous environment with AFM, with a height of 1 nm (Fig S1, in the supplement). Figure 1B shows an AFM image of the surface of bare graphene channel of a graphene-FET; the graphene layer rests on a silicon substrate. Figure 1C shows the same channel after modification with SGP. The difference in height between the samples is 2 nm as expected from the molecular size of SGP (Fig. S1). A high resolution image of the SGP-functionalized surface is presented in Fig. 1D.

**Fig. 1.**
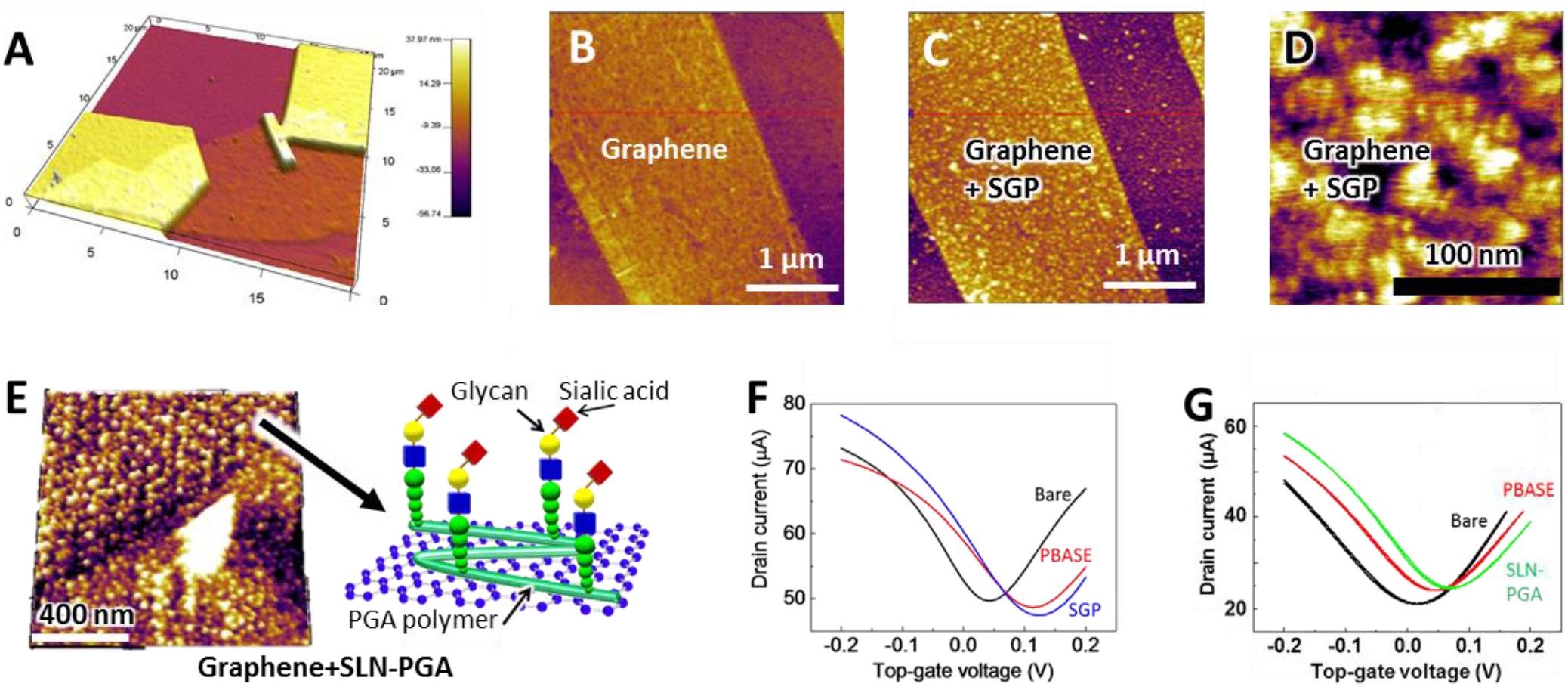
Functionalization of graphene and detection of molecular binding with FET. **(A)** A typical AFM image of graphene on silicon substrate. (**B**) AFM image of bare graphene on the surface of a silicon wafer in water; (**C**) the same graphene surface as in **(B)** after modification with SGP. (**D**) High resolution image of the SGP functionalised surface. (**E**) Graphene surface functionalised with SLN-PGA and cartoon representing the functionalisation scheme. (**F**) Graphene-FET transfer characteristics for bare graphene (black), PBASE-functionalised graphene (red) and SGP-functionalised graphene (blue). (**G**) Graphene-FET transfer characteristics for bare graphene (black), PBASE-functionalised graphene (red) and SLN-PGA-functionalised graphene (green). In all transfer characteristics of the graphene-FET, drain current was switched from hole current to electron current at the charge neutrality point (current minimum) with increasing top-gate voltage.

In order to create a more homogeneous functionalization of the surface of graphene surface where AFM imaging and analysis can be performed more readily, another molecule immobilization method was developed (see Materials and Methods). Briefly, a PBASE-modified graphene surface was exposed to a solution of a sialic acid-containing glycopolymer (SLN-PGA). A cartoon showing the immobilization rationale of SLN-PGA is shown in Fig. 1E. SLN-PGA produces a homogenous functionalization of the surface as seen in Fig. 1E.

Measurements of the drain current of the graphene-FET vs. the top-gate voltage applied from an electrode immersed in solution (transfer characteristics) for all the graphene structures described above were performed in buffer solution. Fig. 1F compares bare graphene, PBASE functionalized graphene and SGP functionalized graphene. All the surfaces present different transfer characteristics. Equivalently, the transfer characteristics for another FET modified with SGP also show a clear shift with respect to bare graphene. Fig. 1G shows results for bare graphene, PBASE- and SLN-PGA-functionalized graphene. Both for SGP and SLN-PGA functionalization the transfer-characteristics of the graphene-FET shifted towards the positive gate voltage direction, as expected from immobilization of negatively charged molecules at the graphene channel (*13*). Both SGP and SLN-PGA have negatively charged sialic acid termini.

### Detection of the binding of HA to the sialic acid molecules at the graphene-FET channel and structural characterisation with AFM

In order to study the binding of HA at the graphene surface, the biomimetic graphene surfaces functionalized with sialic acid were exposed to a solution containing HA. A cartoon showing the surface functionalization strategy is shown in Fig. 2A. Figure 2B shows an AFM image where the three layers present in the surface are exposed. The AFM tip was used to remove the layers by scanning concentric square frames at increasing forces. The deepest surface shows the underlying graphene surface, the second layer represents the surface modified with SLN-PGA and finally the outermost layer shows HA immobilized on SLN-PGA. A histogram of the pixel height distribution of the AFM image is given in Fig. 2C. The average distance between the graphene and the top of the HA layer is 12 ± 3 nm. Figure 2D shows the transfer characteristic of a graphene-FET modified with SLN-PGA before and after exposure to the HA solution (408 nM). The introduction of HA produces a shift towards the negative direction, which reflects the presence of a positively charged molecule (HA) at the channel (*13*). The drain current changes of graphene-FET divided by transconductance correspond to horizontal shift widths of the transfer characteristics. The changes increased with increasing concentration of HA, following Langmuir adsorption isotherm (Fig. 2E). These electrical data also supports the proper structuring of the molecules on graphene biointerface.

**Fig. 2.**
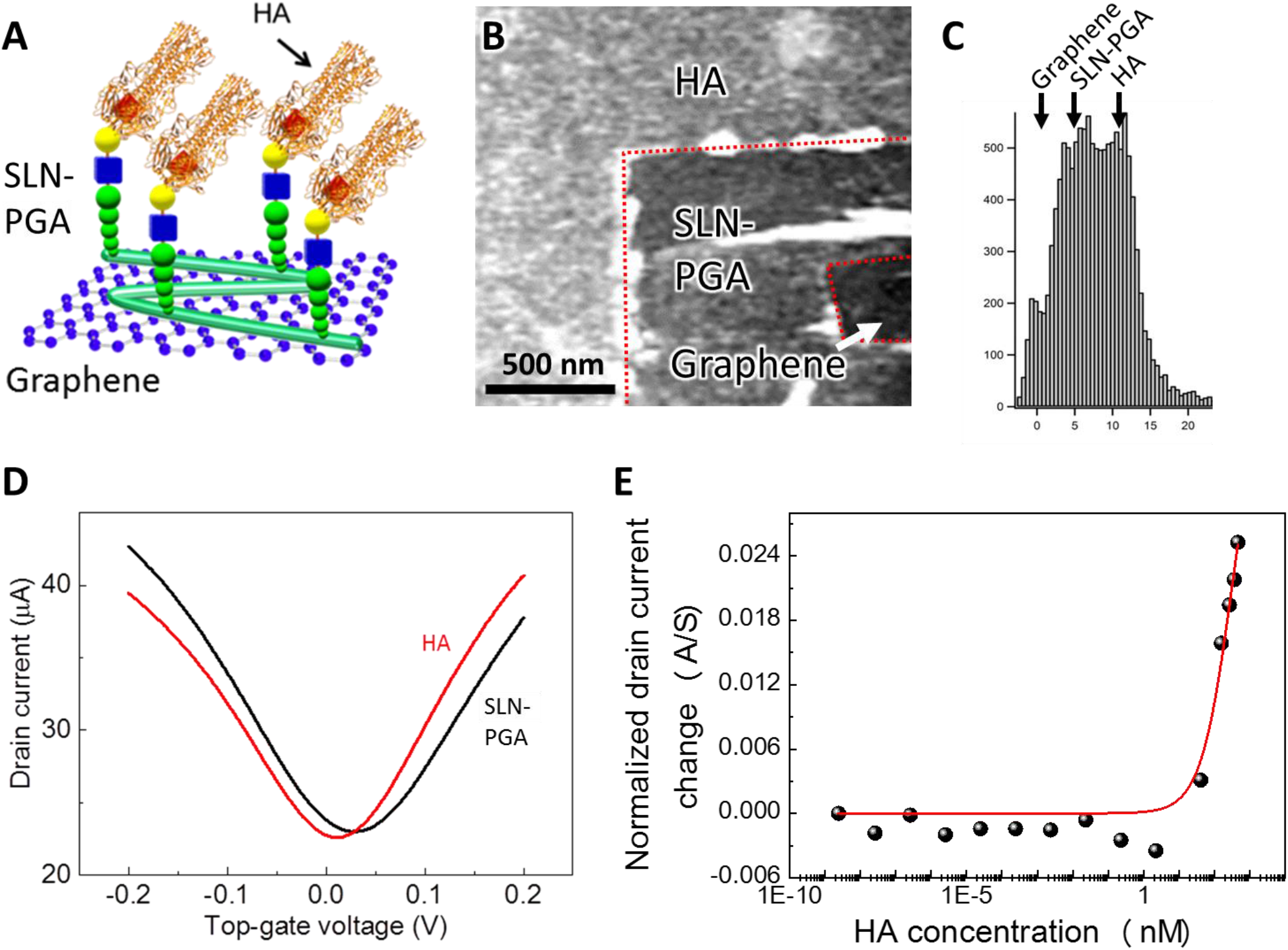
Graphene-FET detection of HA binding. (**A**) Cartoon showing the immobilization of SLN-PGA on graphene and the binging of HA. (**B**) AM-AFM image in solution of the three layers present at the surface: the deepest surface shows the underlying graphene surface, the second layer is the SLN-PGA layer and finally the outermost layer is HA. A histogram of the pixel height distribution of the AFM image is given in (**C**). (**D**) The transfer characteristics of a graphene-FET modified with SLN-PGA before and after exposure to the HA solution. (**E**) Drain current changes of graphene-FET with HA under fixed top-gate voltage at −20 mV. Drain current values were normalized by transconductance of graphene-FET. The data was fitted by Langmuir adsorption isotherm (red curve).

### Imaging the cleavage of the sialic acid groups by the NA enzyme by in-situ AFM

As explained in the introduction, NA enzymes are able to cleave the sialic acid groups *in vivo* and in cell cultures to allow the viruses to escape from the host-cell surface. We exposed a surface where HA is bound to SLN-PGA molecules to NA. A cartoon showing the surface functionalization is given in Fig. 3A. AFM images of the graphene channel on the silicon substrate before and after exposure to NA are shown in Fig. 3B and 3C. The images show the interface of the graphene channel with the underlying silicon. A pixel height distribution for each image is presented in both Fig. 3B and 3C. The average height difference between the silicon and the graphene areas in Fig. 3B is approximately 4 ± 2 nm while the difference between both areas in Fig. 3C is approximately 10 ± 4 nm. This height difference indicates that NA preferentially reacts with the graphene side of the sample, presumably because the HA-immobilized on SLN-PGA facilitates the binding of NA by exposing the right orientation of the protein and the sialic acid, which should not be present on the silicon surface. The broader distribution of the histogram after introduction of NA indicates that NA is probably able to cleave the sialic acid groups from the surface and then accumulate in clumps on top of the HA surface as shown in Fig. 3D and we pictorially propose in Fig. 3E.

**Fig. 3.**
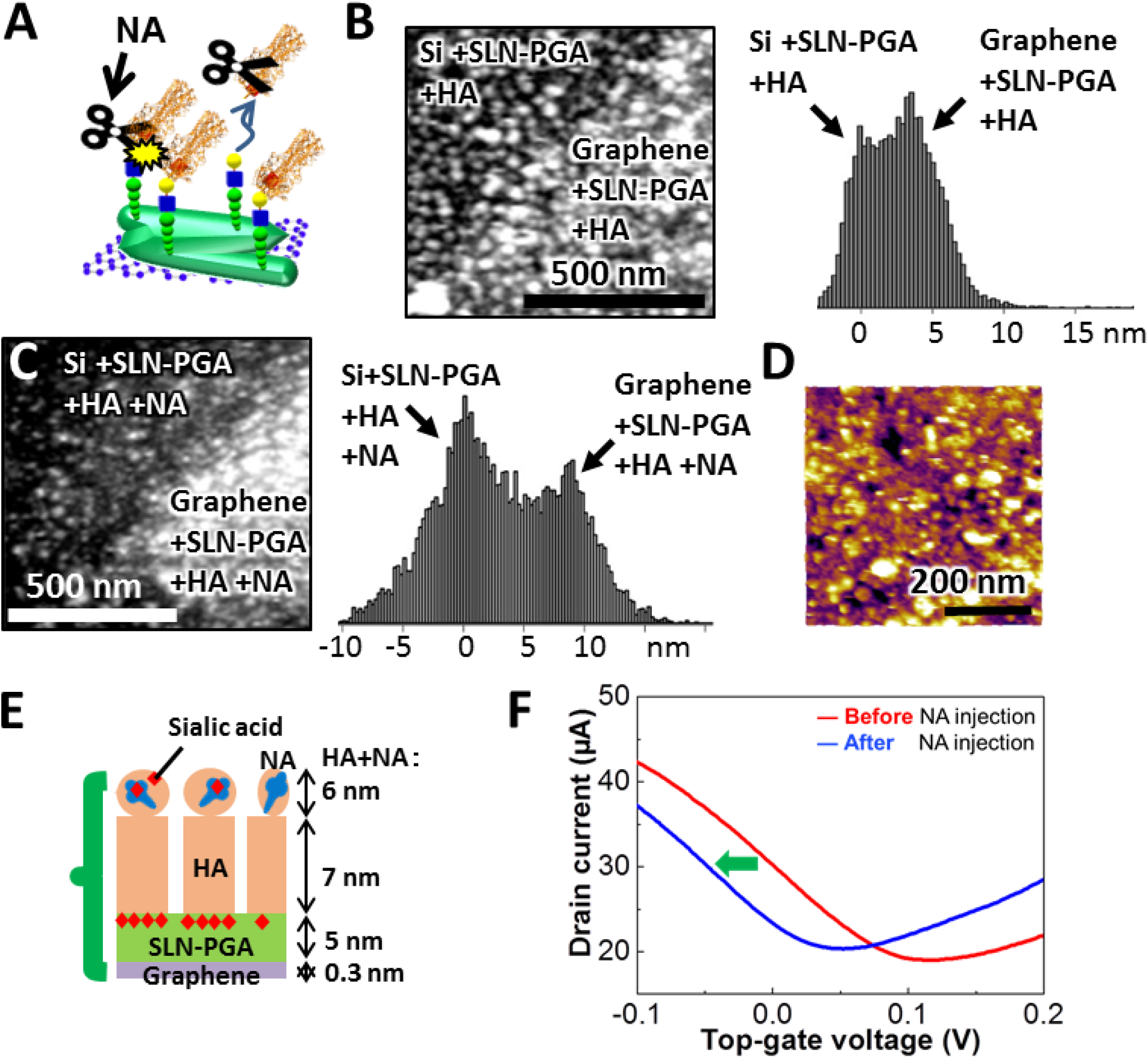
Detecting the cleavage of the sialic acid groups by NA. (**A**) Cartoon showing the surface functionalisation with SLN-PGA and the cleavage of the sialic acid groups by NA. AFM images of the graphene channel on the silicon substrate before and after exposure to NA are shown in (**B**) and (**C**), respectively. Pixel height distribution for each image is presented. (**D**) shows a zoom in area in **c** on the graphene side showing pits indicating that NA is probably able to cleave the bond of sialic acid terminus from the surface and then accumulate in clumps on top of the HA surface, as we propose in the cartoon in (**E**). (**F**) Graphene-FET transfer characteristics measured before and after introduction of NA.

In order to test the effect of NA, we used a graphene-FET set up exposing a SGP-covered graphene surface. A solution containing NA was introduced to the graphene-FET channel, and the transfer characteristics were measured before and after introduction of NA as shown in Fig. 3F. The transfer characteristics shift towards the negative voltage direction after NA injection. This indicates that NA is cleaving the bond of negatively-charged sialic acid terminus, as a negative shift can be associated with a more positively-charged surface.

### Effect of antiviral zanamivir drugs on NA activity: AFM imaging and FET monitoring

In order to investigate the effect of zanamivir on the interaction of NA with HA-SLN-PGA complex with AFM, we first prepared a sample of graphene-FET with HA immobilized on its surface. After that the surface immersed in a solution containing the antiviral drug zanamivir. Subsequently we exposed the surface to the NA/zanamivir mixture solution. A cartoon showing the surface functionalisation of graphene and the hypothesis of the experiment is given in Fig. 4A.

**Fig. 4.**
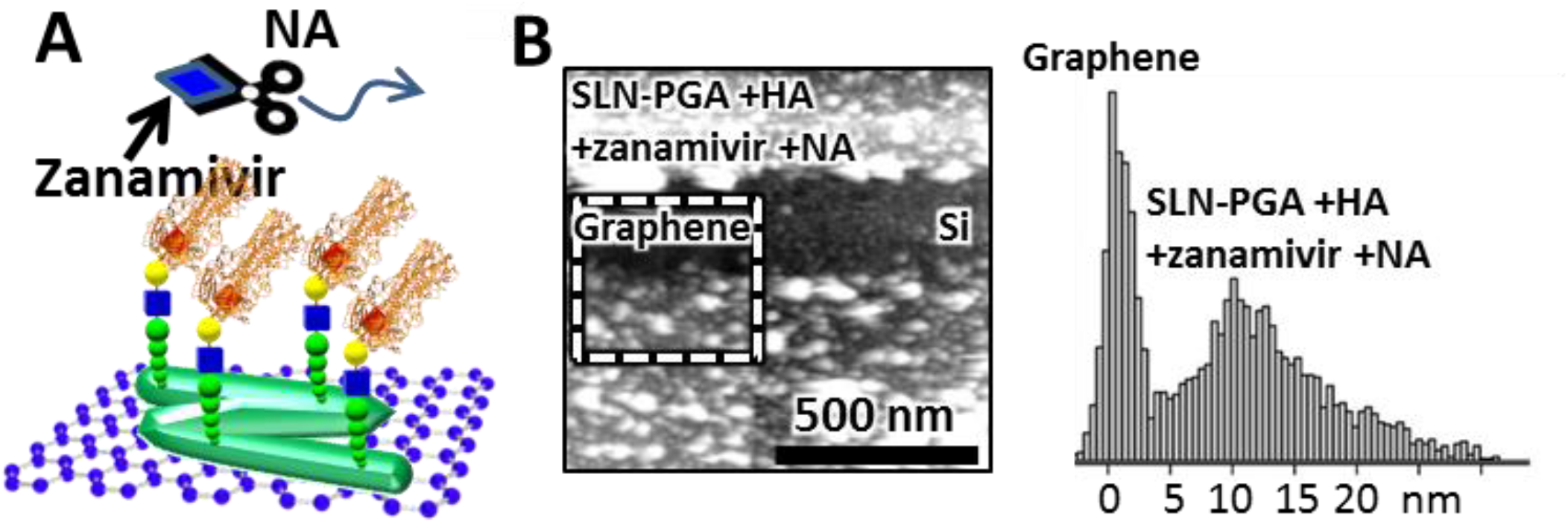
Effect of zanamivir on the binding of NA to the protein layer. (**A**) Cartoon showing the hypothesis of the experiment with HA immobilized on the SLN-PGA functionalising the graphene. (**B**) AFM image in solution showing the bare graphene surface and the effect of flowing NA on the surface functionalised with HA as shown in (**A**). The pixel size distribution corresponding to the area confined by the white dotted square is given to the right of the AFM image.

Measurement of the heights of each of the biomolecular layers was made in an area of the sample where the AFM was used to strip the protein coating from the surface. The AFM image shows the protein layer, the bare graphene and the underlying silicon surface as shown in Fig. 4B. A histogram of the pixel height obtained from a portion of the image corresponding to the graphene and the SLN-PGA+HA+zanamivir+NA layer (white dotted square) is also given in Fig. 4B. The height difference is 12 ± 8 nm that is equivalent to the height obtained for the SLN-PGA+HA images shown in Fig. 2B. This indicates that NA could not react to the protein layer in the presence of zanamivir.

In order to monitor, in-situ, the effect of the injection of NA on the graphene-FET where SGP molecules had been immobilized on the surface, the hole current of the graphene-FET was monitored in real-time. Fig. 5A shows the resulting behaviour with a top-gate voltage of −50 mV, where NA was introduced at time zero. After Introduction of NA, the hole current decreased exponentially, likely due to the release of negatively-charged sialic acid from the graphene surface (*13*), the experimental exponent obtained from the fitting is 1.72×10^−3^ s^−1^ which corresponds to the reaction rate.

**Fig. 5.**
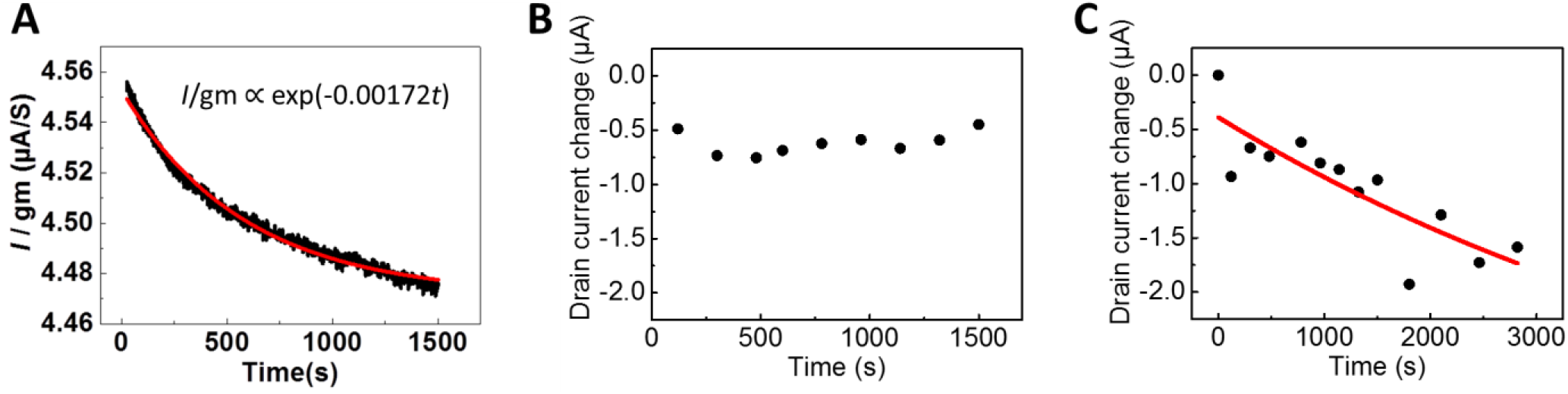
In situ, real time monitoring the effect of NA on SGP functionalised graphene-FET. (**A**) Hole current vs. time with a top-gate voltage of −50 mV, where NA was introduced at time zero. The current decreases indicating cleavage of the sialic acid contained in SGP by NA. In (**B**) the effect of zanamivir on NA was studied using the hole current vs. time graph, the current remains constant indicating that zanamivir inhibits the binding of NA to the SGP molecule. The NA reaction was recovered after zanamivir removal from the graphene-FET as shown in (**C**).

To investigate the validity of this measurement we estimate the expected value of the exponent using Michaelis–Menten reaction kinetics theory. In this model, NA reactions should follow the equation:

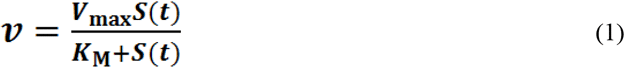

where *v* is the reaction rate, and *V*_max_, *K*_M_ and *S*(*t*) are maximum reaction rate, Michaelis constant and substrate concentration as a function of time, respectively. *V*_max_ was determined as 1.7 s^−1^ by conventional colorimetric assay. *K*_M_ value of NA reported in literatures are in the range of 10^−4^ M (*19, 20*). In our reaction system, SGP is confined to the surface of graphene, and hence *S*(*t*) << *K*_M_ (See Fig. S2 and supplementary material discussion for assumptions used to make this approximation). Therefore, equation (1) can be approximated by:

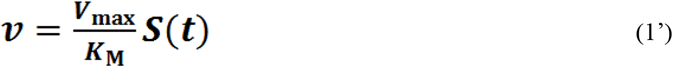

From equation (1),

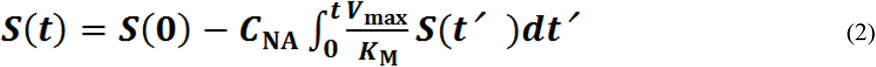

where *C*_NA_ is NA concentration (200 nM). *S*(*t*) shows exponential decay and the time constant *τ* is:

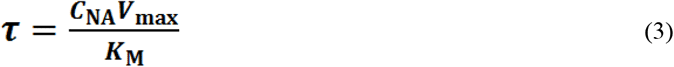

This time constant obtained from curve fitting (1.72×10^−3^ s^−1^) was well consistent with the calculated *τ*, as *K*_M_ value obtained from the fitting was 2×10^−4^ M. Therefore we can conclude that the graphene-FET quantitatively and accurately monitored the NA reaction on the two-dimensional biomimetic graphene surface in real time.

Finally the effect of zanamivir on the hole current is shown in Fig. 5B. First zanamivir was introduced on a graphene-FET exposing a SGP layer immobilized on the graphene. Zanamivir was also mixed with NA outside of the graphene-FET. NA was then introduced to graphene-FET with monitoring the drain current. As shown in Fig. 4B, even after NA introduction the current remains constant indicating that NA is not reacting to the sialic acid in the presence of zanamivir. After zanamivir removal, the drain current started to decrease, indicating that the interaction at the graphene surface was recovered (Fig. 5C). Although the time constant of the reaction was not perfectly recovered its initial value obtained by conventional colorimetric assay, probably due to residual zanamivir at the graphene interface, these results show the potential of the lab-on-a-graphene-FET for the evaluation of drug candidates in pharmaceutical research.

## Conclusions

We have described the fabrication of a two-dimensional structure that mimics the surface of the host cell, constructed using graphene as a template to immobilize key proteins related to viral infection. We demonstrated that it is possible to create a surface presenting sialic acid that is useful for on-chip studies of the interactions of HA and NA enzymes of the influenza virus. Our results demonstrate that graphene is a suitable material to study molecular interactions that happen at surfaces by high resolution microscopies, in this case in situ AFM imaging. Our AFM experiments confirm previous results of the molecular interactions (*2*), namely that HA binds to the sialic acid groups at the surface and that NA is able to cleave the sialic acid groups.

Furthermore we demonstrate that cell surface biomimetic graphene can be successfully used as the channel of a graphene-FET device. The device can then be used as “lab-on-a-graphene-FET” structure able to monitor viral protein molecular binding events and the effect of drugs on the key interactions, in-situ and in real time. Our graphene-FET device is able to detect binding of HA to the sialic acid groups, and the cleavage of the sialic acid bond by NA. Furthermore the time course of the interaction of NA to sialic acid groups was measured, from which the correct reaction rate was calculated. The inhibitory effect of zanamivir on the NA-sialic acid interactions was also detected and monitored with the FET.

Taken together our results demonstrate the potential of graphene and graphene-FETs for the study of molecular interactions at surfaces and in biomedical assays and drug discovery.

## Materials and Methods

### Graphene-FET preparation and measurements

Freshly exfoliated graphene was deposited at a Si/SiO_2_ substrate surface and Au/Ni electrodes were fabricated through electron-beam lithography, electron-beam deposition and lift-off process. Then the graphene-FET was annealed at 300 °C for 30 minutes in Ar/H_2_ atmosphere to remove contamination on graphene surface.

### Molecular functionalisation of the graphene surface

The graphene-FET was immersed in 2-methoxyethanol solution containing 1 mM PBASE (Thermo Fisher Scientific Inc.) for 1 hour. During this process, pyrenyl group of PBASE formed strong π-π stacking with graphene (*17*). Immediately after the immersion, the graphene-FET was rinsed in pure water and again immersed in 1 μM α2,6-SGP (Fushimi Pharmaceutical Co., Ltd., Japan) aqueous solution overnight. PBASE functions as a linker molecule between the surface graphene and SGP, owing to a succinimide group forming covalent bond to N-terminus of SGP.

For AFM imaging experiments, the PBASE-modified graphene surface was immersed for 18 hours in a 100 nM of the sialic acid-containing glycopolymer (Neu5Acα2-6Galβ1-4GlcNAcβ-poly γ-L-glutamic acid, SLN-PGA) in phosphate buffer solution (10 mM, pH 6.0). The synthesis of SLN-PGA is described elsewhere (*21*). In SLN-PGA, 52.2% of side chains in poly γ-L-glutamic acid were replaced by sialoglycan. The average molecular weight of SLN-PGA is 3,528 kDa. During this process the succinimide group of PBASE makes a covalent bond with amide group of the SLN-PGA.

Binding of HA to SLN-PGA was studied by immersion of the SLN-PGA-graphene sample in a 300 nM HA derived from Influenza H1N1 (A/California/07/2009), recombinant from human embryonic kidney (HEK) 293 cells (Sino Biological Inc., China) solution in phosphate buffer solution (10 mM, pH 6.0).

The effect of NA on the HA-SLN-PGA-graphene sample was studied upon immersion of the sample in 400 nM NA from *Arthrobacter ureafaciens* (Nacalai Tesque, Inc., Japan) solution in phosphate buffer (10 mM, pH 6.0) for 15 minutes, after which NA cleaved the sialic acid terminal of SLN-PGA.

Finally 133 nM of NA was introduced after adding 50 mM zanamivir to the solution (Abcam plc., UK).

### FET measurements

A silicone reservoir was placed on the graphene-FET to store the assay liquid solution. The assay buffer was 10 mM sodium phosphate (pH 6.2). All electrical measurements were carried out using a semiconductor parameter analyzer (Keysight Technologies B1500A). An Ag/AgCl reference electrode was used to apply top-gate voltage to the graphene channel through the assay liquid solution. The drain voltage was fixed at 0.1 V in all measurements. For electrical inhibition assay viral NA derived from influenza H1N1 (A/California/04/2009), recombinant from HEK 293 cells (Sino Biological Inc., China) was used (Fig. 5B and 5C).

### AFM measurements

AFM imaging was carried out in an MFP-3D (Oxford Instruments Asylum Research, Santa Barbara, CA.). Experiments were done with TR800 cantilevers (Olympus, Japan) with spring constant 0.7 N/m using low amplitude amplitude-modulation (AM) AFM. AFM was used to study of (i) unmodified pure graphene, (ii) immobilized SLN-PGA on graphene (SLN-PGA-Graphene), (iii) HA bound to SLN-PGA on graphene (HA-SLN-PGA-Graphene), (iv) NA on HA-SLN-PGA-Graphene samples (NA-HA-SLN-PGA-graphene), and (v) zanamivir effect on NA with HA-SLN-PGA-Graphene (Zanamivir-NA-HA-SLN-PGA-graphene).

All measurements were performed in phosphate buffer at 10 mM, pH 6.0. After each step the graphene electrode was gently rinsed with phosphate buffer. In some cases the AFM was used at high (>10 nN) scanning forces to selectively remove molecular layers from the surface.

Images were flattened using Asylum Research software based on Igor Pro (Wavemetrics Inc. Oregon, USA), the height histograms were fitted to Gaussian functions using Igor Pro to find the average distance between the peaks to determine the relative height of the molecular layers.

## Acknowledgments

KM, SC, KK, RH acknowledge support from the Japan Society for the Promotion of Science (JSPS) Core-to-Core grant, ARP acknowledges support from Engineering and Physical Sciences Research Council. TO acknowledges supports from JSPS KAKENHI (16K13638 and 18K14107). This study was also supported by JST-PRESTO (JPMJPR19G3), CREST (JPMJCR15F4) and Mirai (JPMJMI19D4). We are grateful to the Yamasa Corporation for the generous gift of the multivalent sialyloligosaccharides with a γ-polyglutamic acid backbone.

TO, KK RH did current measurement experiments with FETs, KK, RH, ARP, CG, SC did AFM experiments, YS, SN synthesized the molecules for functionalization of graphene. SC, TO, KK wrote the paper. All author contributed to experimental design and discussed the results. All authors read and approved the text.

The authors declare that they have no conflict of interest.

