## Supplementary Materials for "Lab-on-a-graphene-FET detection of key molecular events underpinning influenza virus infection and effect of antiviral drugs"

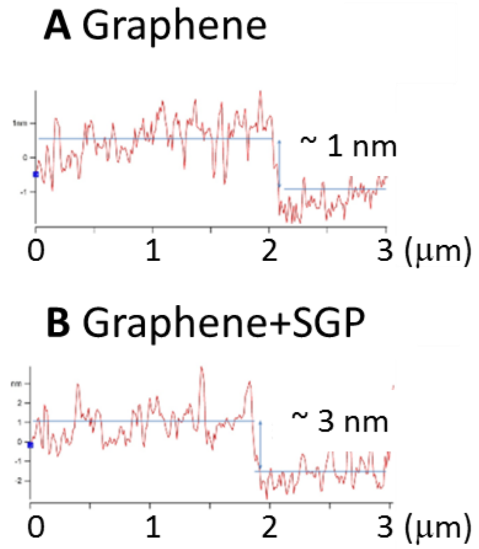


**Fig. S1:** Comparison of average height of graphene on the silicon substrate measured in Fig. 1B and of graphene functionalised with SGP corresponding to Fig. 1C. Graphene thickness in (**A**) was increased probably due to hydration.

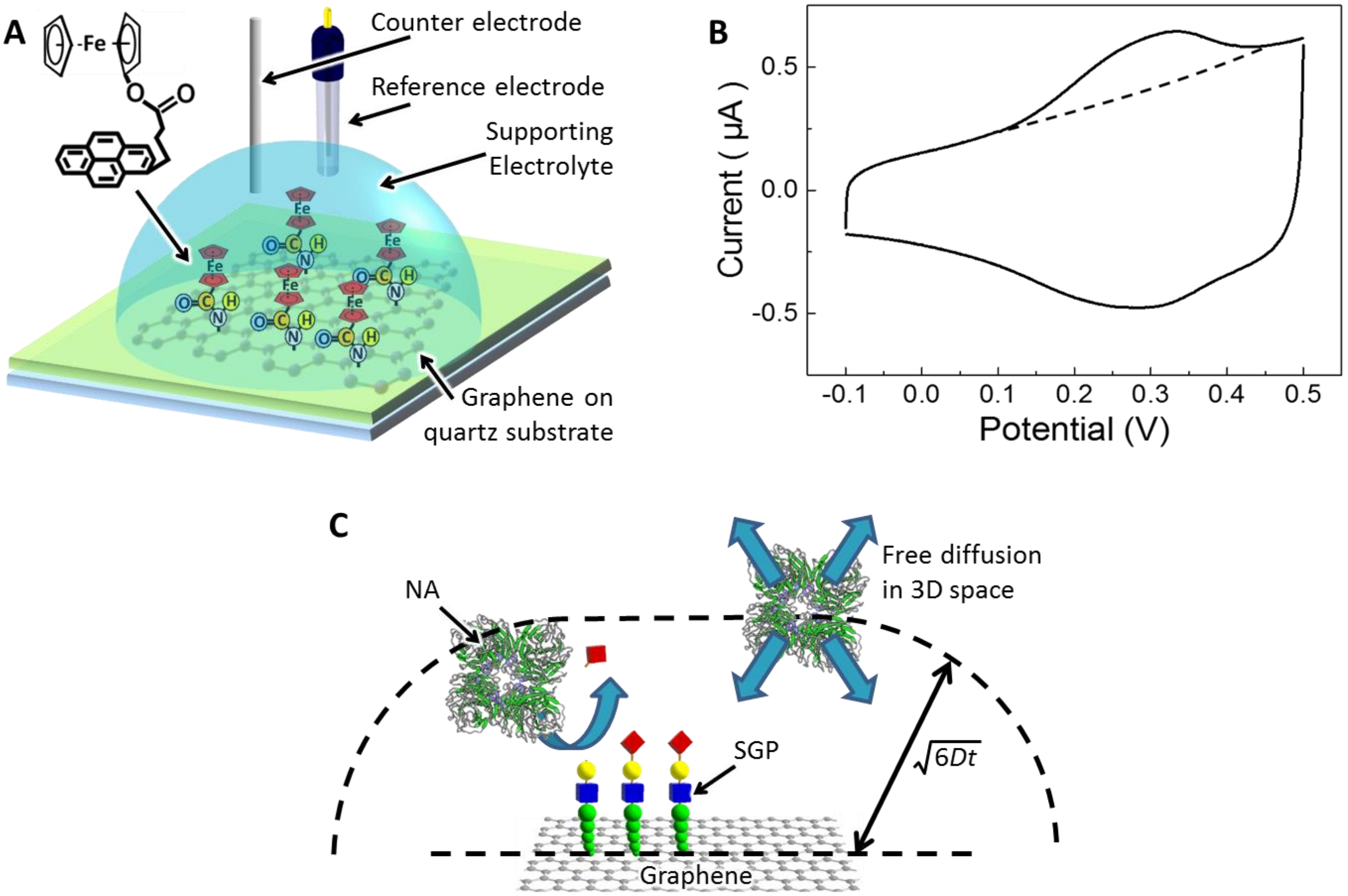


**Fig. S2:** Neuraminidase reaction for substrates fixed on the graphene surface. (**A** and **B**) Estimation of immobilization density of pyrenyl linkers on graphene surface. (**A**) Schematics of cyclic voltammetry experiment. Ferrocene-binding PBASE linker was immobilized to graphene grown by chemical vapor deposition of 0.5 $cm^2$ area. A platinum electrode, an $Ag/AgNO_3$ electrode and tetrabutylammonium hexafluorophosphate in dichloromethane (0.1 M) were used as a counter electrode, a reference electrode and supporting electrolyte, respectively. (**B**) Cyclic voltammogram for ferrocene-immobilized graphene electrode. The sweep speed was 0.1 V/s. From the oxidation peak area above the dotted line, the surface density of ferrocene on graphene was estimated as 13 $pmol/cm^2$. The channel size of graphene-FET was typically 5×4 μm. Therefore, the amount of sialic acid on a graphene channel is considered to be $3.2\times10^6$ molecules or less (note that one SGP molecule has two sialic acid termini). (**C**) Model to calculate the reaction rate on graphene surface. Only molecules reaching the surface contribute to the reaction. The diffusion coefficient *D* was estimated from molecular weight of NA, 52~88 kDa, as 56~67 $\mu m^2/s$ (*22*). Therefore, within 0.6 s after reaction started, effective reaction volume increased, so that the concentration of sialic acid fixed on the graphene reached less than 1% of $K_M$. It indicates $S(t) << K_M$ in the experimental timescale of this study.
